# Independent duplications of the Golgi phosphoprotein 3 oncogene in birds

**DOI:** 10.1101/2021.01.31.429031

**Authors:** Juan C. Opazo, Michael W. Vandewege, Javier Gutierrez, Kattina Zavala, Luis Vargas-Chacoff, Francisco J. Morera, Gonzalo A. Mardones

**Affiliations:** Integrative Biology Group, Universidad Austral de Chile, Valdivia, Chile; Instituto de Ciencias Ambientales y Evolutivas, Facultad de Ciencias, Universidad Austral de Chile, Valdivia, Chile; Millennium Nucleus of Ion Channel-Associated Diseases (MiNICAD), Valdivia, Chile; Department of Biology, Eastern New Mexico University, Portales, New Mexico, USA; Instituto de Ciencias Marinas y Limnológicas, Universidad Austral de Chile, Valdivia, Chile; Centro Fondap de Investigación de Altas Latitudes (IDEAL), Universidad Austral de Chile, Valdivia, Chile; Applied Biochemistry Laboratory, Instituto de Farmacología y Morfofisiología, Facultad de Ciencias Veterinarias, Universidad Austral de Chile, Valdivia, Chile; Department of Physiology, School of Medicine, Universidad Austral de Chile, Valdivia, Chile; Center for Interdisciplinary Studies of the Nervous System (CISNe), Universidad Austral de Chile, Valdivia, Chile

**Keywords:** Birds, Cancer, Gene Family Evolution, Golgi apparatus, GOLPH3, GOLPH3L, intrinsically disordered region, Vps74

## Abstract

Golgi phosphoprotein 3 (GOLPH3) was the first reported oncoprotein of the Golgi apparatus. It was identified as an evolutionarily conserved protein upon its discovery about 20 years ago, but its function remains puzzling in normal and cancer cells. The GOLPH3 gene is part of a group of genes that also includes the GOLPH3L gene. Because cancer has deep roots in multicellular evolution, studying the evolution of the GOLPH3 gene family in non-model species represents an opportunity to identify new model systems that could help better understand the biology behind this group of genes. The main goal of this study is to explore the evolution of the GOLPH3 gene family in birds as a starting point to understand the evolutionary history of this oncoprotein. We identified a repertoire of three GOLPH3 genes in birds. We found duplicated copies of the GOLPH3 gene in all main groups of birds other than paleognaths, and a single copy of the GOLPH3L gene. We suggest there were at least three independent origins for GOLPH3 duplicates. Amino acid divergence estimates show that most of the variation is located in the N-terminal region of the protein. Our transcript abundance estimations show that one paralog is highly and ubiquitously expressed, and the others were variable. Our results are an example of the significance of understanding the evolution of the GOLPH3 gene family, especially for unraveling its structural and functional attributes.

## Introduction

Golgi phosphoprotein 3 (GOLPH3) is a highly conserved protein of the Golgi apparatus ^1,2^ considered the first oncoprotein of this subcellular compartment ^3^. The GOLPH3 gene family comprises the conserved GOLPH3 gene and the GOLPH3L gene found only in vertebrates ^1,2^. Despite the vast amount of empirical evidence demonstrating the contribution of GOLPH3 to tumorigenesis and cancer, a full understanding of its molecular role has not yet emerged. This is mainly due to multiple functions attributed to GOLPH3 ^3^, including, the sorting of Golgi glycosyltransferases ^4–6^, the modulation of focal adhesion dynamics ^7^, induction of membrane curvature ^8^, and an intriguing function for a Golgi protein, regulating mitochondrial function ^9^. Because cancer has deep evolutionary roots that arise as a consequence of the multicellularity ^10^, and is widespread across animals ^11^, studying the evolution of the GOLPH3 gene family in non-model species can provide significant information for a comparative oncology approach, which is emerging as an integrative field to tackle cancer ^10^.

The availability of whole-genome sequences opens an opportunity to understand the evolution of gene families. The annotation of gene repertoires in different species has revealed that copy number variation is an important source of variability that should be considered when making functional comparisons ^12,13^. Phylogenetic reconstructions show that the evolution of gene families follows complex pathways, including gene gain and losses and independent origins, making it challenging to perform direct interspecies comparisons. Thus, understanding the variability of gene repertoires and their duplicative history represents an essential piece of information to understand the biological functions associated with a group of genes and make biologically meaningful comparisons. Today, the GOLPH3 gene family is viewed as a group of genes containing two paralogs (GOLPH3 and GOLPH3L) with 1:1 orthologs among most vertebrate species ^1,2^. Among amniotes, it is suggested that, in contrast to mammals, birds are less susceptible to cancer^10,14–18^; however, this information should be taken with caution given sampling bias ^19^. Thus, the study of genes associated with cancer in birds could provide clues about the genetic bases associated with this difference and suggest additional model systems that could help to understand the biology of the GOLPH3 gene family.

The main goal of this study was to analyze the evolutionary history of the GOLPH3 gene family in birds. We took advantage of whole genome sequences in representative species of all main lineages of birds to understand the evolutionary pathways that gave rise to GOLPH3 paralogs. According to our assessment, we identified a repertoire of three GOLPH3 genes in birds. We found duplicated copies of the GOLPH3 gene in all main groups of birds other than paleognaths, and a single copy of the GOLPH3L gene that would be derived from the common ancestor of all birds. In the case of the GOLPH3 gene, our gene tree suggests at least three independent origins for the duplicated copies, in the ancestor of Galliformes and Anseriformes, in the ancestor of Anseriformes, and the ancestor of Neoaves. Divergence estimates between duplicated genes showed that most of the variation is located in the N-terminal region of the protein. Our transcript abundance estimations showed that one paralog was highly and ubiquitously expressed, while the others were variable. Our evolutionary analyses suggest a more complex than anticipated evolutionary history of the GOLPH3 gene family, a scenario that could have implications for cancer.

## Results and Discussion

### Independent duplication events characterize the evolution of the GOLPH3 gene family member in birds

Comparing the sister group relationship among gene family members, i.e. gene tree, with the species tree represents a fundamental strategy to understand homologous relationships, duplicative history, and modes of evolution of any group of genes ^20,21^. In our case, our gene tree did not significantly deviate from the most updated phylogenetic hypotheses for the main group of birds ^22–25^(Fig. 1), suggesting that GOLPH3 was present in the ancestor of the group as a single copy gene. We recovered a clade containing GOLPH3 sequences from paleognaths (ostriches, tinamous, and allies) sister to GOLPH3 sequences from all other birds (Fig. 1). Further, we recovered the sister group relationship of the GOLPH3 sequences from Galliformes (chickens, pheasants, and allies) and Anseriformes (ducks, swans, and allies), in turn, this clade was recovered sister to GOLPH3 sequences from Neoaves (Fig. 1).

**Figure 1.**
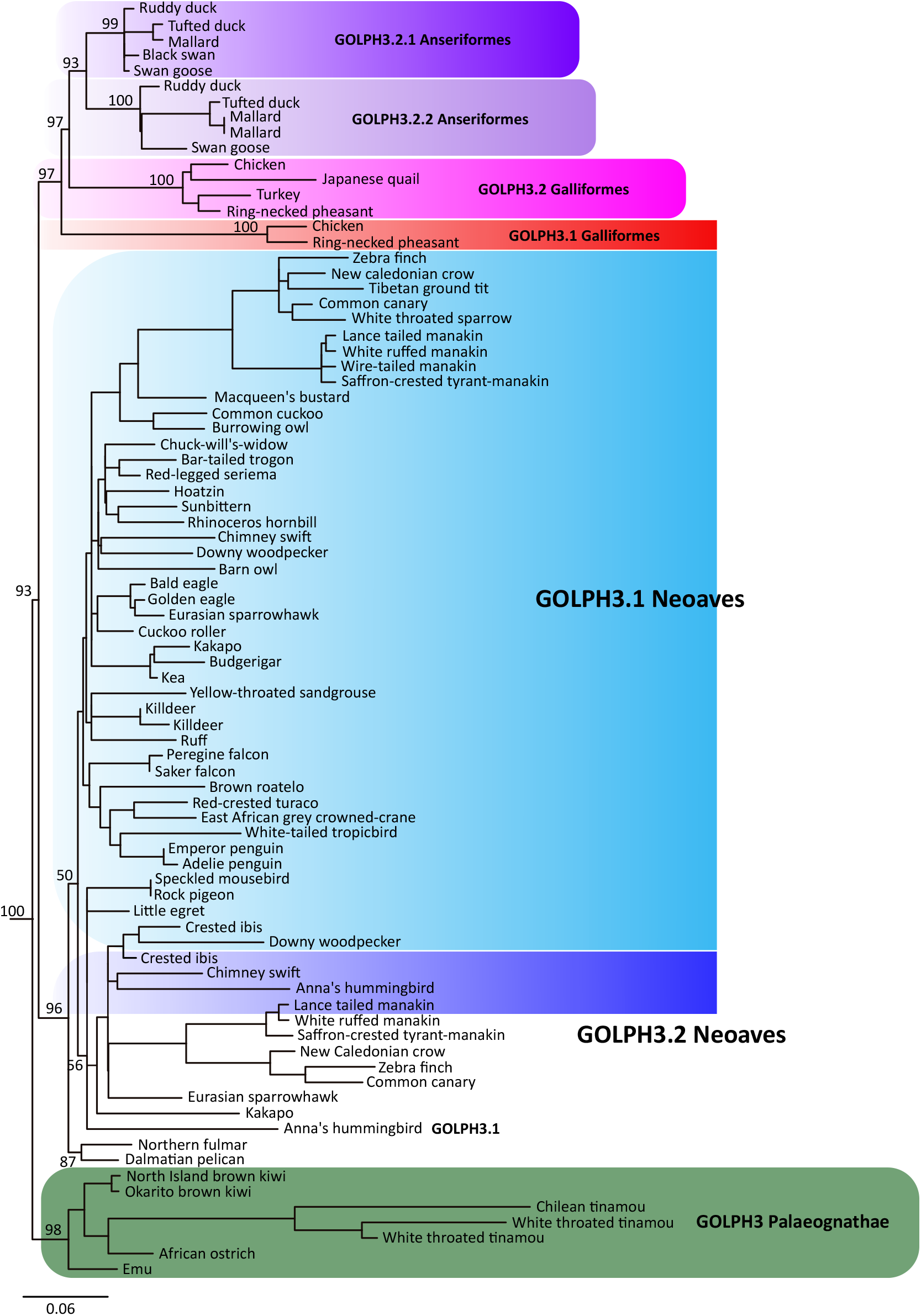
Maximum likelihood tree showing sister group relationships among GOLPH3 genes of birds. Numbers above the nodes correspond to support values from the ultrafast bootstrap routine. GOLPH3 sequences from crocodiles and turtles were used as outgroups (not shown). The scale denotes substitutions per site and colors represent gene lineages. In the case of the Northern fulmar and Dalmatian pelican, there is no syntenic information to define to which gene lineage they belong. In consequence, they were not included in any shading or do not have a gene name

We found a single copy gene, located on chromosome Z, in most paleognaths species, except in the white-throated tinamou (*Tinamus guttatus*), where two copies were identified (Fig. 1), suggesting that this species independently gave rise to a second GOLPH3 copy located on chromosome W. The location of these genes on sexual chromosomes, and given the sex-determination system of birds ^26^, indicates that only females (ZW) can express both paralogs. In the case of Neoaves, our tree topology is not well resolved, being difficult to anticipate details regarding the duplicative history of the GOLPH3 paralog in this group (Fig. 1). However, for a diversity of species (e.g., zebra finch, common canary, kakapo), we found duplicated copies on different chromosomes, suggesting that the duplication event that gave rise to them occurred in the ancestor of Neoaves. Similar to the case of the white-throated tinamou (*Tinamus guttatus*), we found duplicated copies in the killdeer (*Charadrius vociferus*) that were recovered sister to each other (Fig. 1), suggesting that they arose as a product of a species-specific gene duplication event.

The evolutionary history of the GOLPH3 gene in the clade that includes Galliformes and Anseriformes followed a more complicated evolutionary pathway (Fig. 1). According to our assessment, we found a repertoire of two copies in species belonging to both groups (Fig. 1); however, our gene tree suggests that the events that gave rise to them followed a pattern of gene birth-and-death ^27^(Fig. 2). The reconciliation of the gene tree with the species tree suggests that the last common ancestor of Anseriformes and Galliformes, which lived 80 million years ago approximately ^28^, had a single copy gene that underwent a duplication event (Fig. 2), giving rise to a repertoire of two GOLPH3 copies (Fig. 2). One of the copies was retained in Galliformes (Fig. 2; GOLPH3.1_GA_; red lineage), but lost in Anseriformes (Fig. 2; red lineage). This GOLPH3 gene copy is located on chromosome W. The other copy, also originated in the ancestor of Galliformes and Anseriformes, was also retained in Galliformes (Fig. 3; GOLPH3.2_GA_; pink lineage) and is located on chromosome Z. In the last common ancestor of Anseriformes, this copy underwent a duplication event giving rise to two copies (Fig. 3; GOLPH3.2.1_A_, purple lineage and GOLPH3.2.2_A_, light purple lineage). Like other cases, in Anseriformes, one of the copies is located on chromosome Z, while the other on chromosome W; the exception is the mallard (*Anas platyrhynchos*) in which is found on chromosome 22.

**Figure 2.**
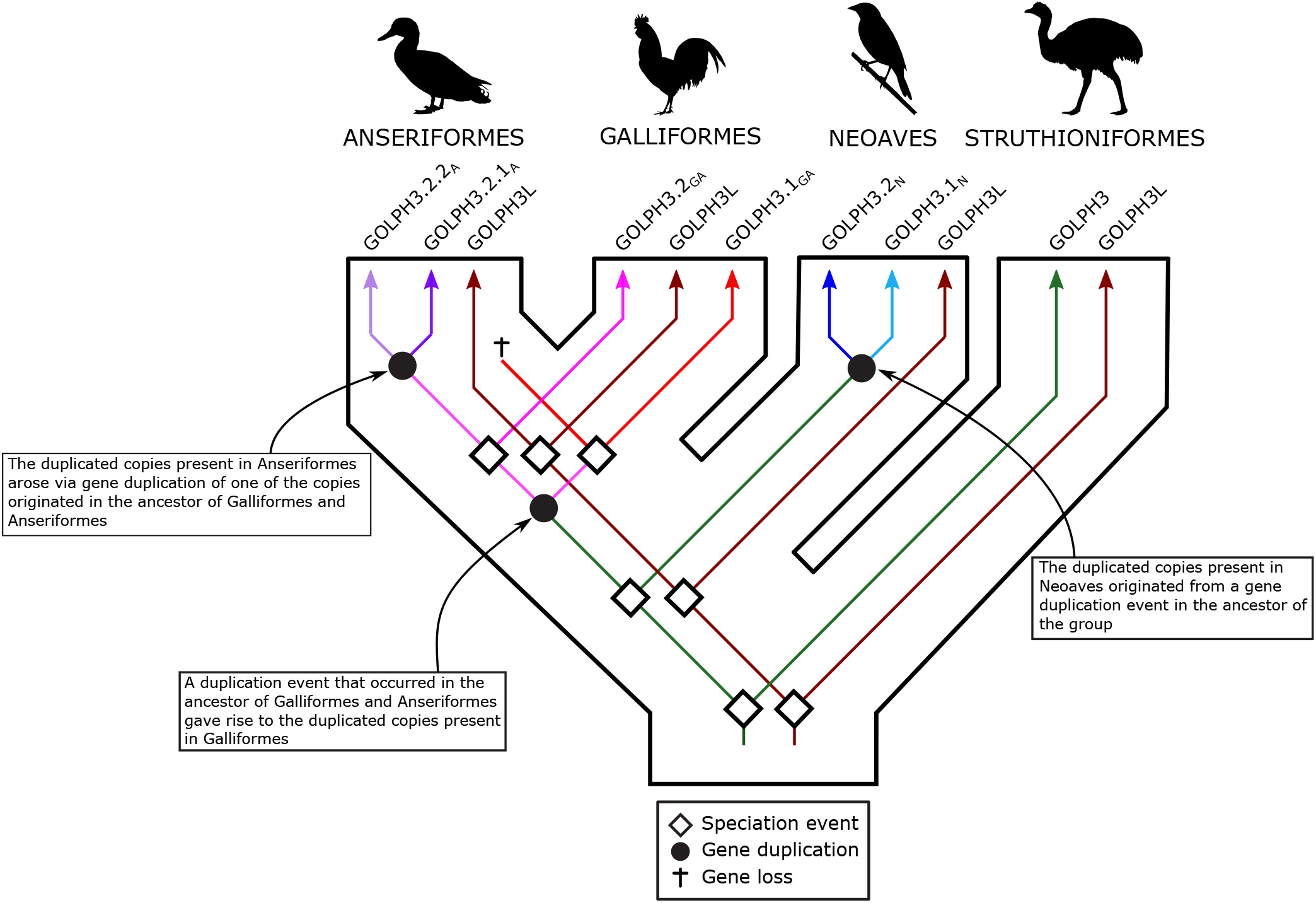
An evolutionary hypothesis regarding the evolution of the Golgi phosphoprotein 3 (GOLPH3) gene family in birds. According to our results, the last common ancestor of birds had a repertoire of two GOLPH3 genes (GOLPH3 and GOLPH3L). The GOLPH3L gene was inherited by all species of birds and maintained as a single copy gene (brown lineage). The evolution of the GOLPH3 gene is more complicated. The gene present in the bird ancestor was inherited by paleognaths (ostriches, tinamous and allies) and maintained as a single copy gene in most species. In the case of Neoaves (zebra finches, kakapos and allies), the ancestral GOLPH3 gene underwent a duplication event in the ancestor of the group giving rise to a repertoire of two genes (blue and light blue gene lineages). In the last common ancestor of Anseriformes (ducks, swans and allies) and Galliformes (chickens, pheasants and allies) the ancestral GOLPH3 gene underwent a duplication event giving rise to a repertoire of two genes, one of the copies was retained in Galliformes (red lineage), but lost in Anseriformes (red lineage). The other copy originated in the ancestor of Galliformes and Anseriformes, was also retained in Galliformes (pink lineage), however, in the last common ancestor of Anseriformes underwent a duplication event giving rise to the repertoire of two copies (purple and light purple lineages). We use the subscripts to indicate the ancestor in which the gene repertoires were originated, N = the ancestor of Neoaves, A = the ancestor of Anseriformes and GA = the galliform/anseriform ancestor. Silhouette images were obtained from PhyloPic (http://phylopic.org/).

**Figure 3.**
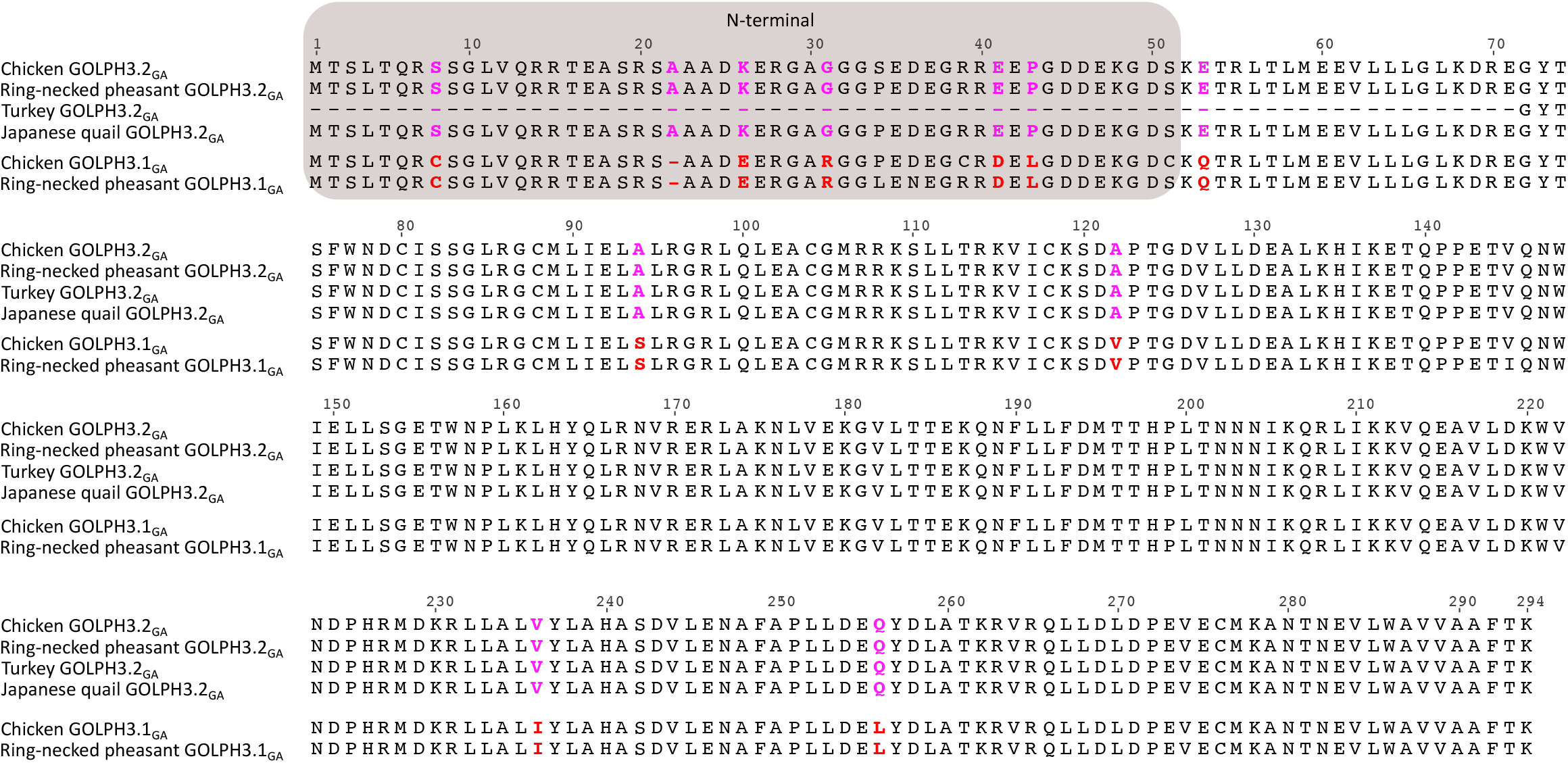
Alignment of Golgi phosphoprotein 3 (GOLPH3) amino acid sequences from chicken (*Gallus gallus*), ring-necked pheasant (*Phasianus colchicus*), turkey (*Meleagris gallopavo*) and japanese quail (*Coturnix japonica*). Amino acids positions that differentiate between paralogs are in pink and red. The N-terminal region of the protein is shaded.

Thus, the main groups of birds possess gene repertoires with different evolutionary origins (Fig. 2). Most paleognaths retained the ancestral condition of a single gene copy, whereas Galliformes, Anseriformes, and Neoaves possess duplicated copies that originated independently (Fig. 2). Anseriformes and Neoaves gave rise to their repertoire in the ancestor of each group (Fig. 2), while Galliformes retained copies that originated in the ancestor of Galliformes and Anseriformes (Fig. 2). The independent origin of gene families in different groups is not an unusual event during the evolutionary process ^29–33^; however, it should be taken into account when making comparisons because non-orthologous genes - i.e., genes with different evolutionary origin - are being compared. Our results also highlight the importance of manual curation in defining the composition of gene families. The description of new genes is also not uncommon ^34–36^, and their discovery could be attributed to their presence in non-model species and/or the absence of appropriate evolutionary analyses. The presence of species with different gene repertoires represents an opportunity to understand the evolutionary fate of duplicated genes ^37^ and the biological functions associated with a group of genes. This phenomenon, variation in gene copy number, has been associated with differences in susceptibility to diseases in different taxonomic groups. For example, in the African elephant (*Loxodonta africana*), it has been claimed that an expansion in the number of TP53 “the guardian of the genome” gene copies could help to explain the lower risk of developing cancer in this large and long-lived animal ^12,13^. Similarly, in whales there are also an expansion of gene families related to cancer, and an accelerated rate of evolution in genomic regions enriched with pathways involved in cancer ^38,39^. Further evidence comes from bats, a group in which the lifespan exceeds the expectation based on their body size ^40^. In this group, it has been documented the expansion of several genes, for example, FBXO31, which is related to cell cycle arrest and response to DNA damage diminishing the probability of developing cancer ^41–43^. Thus, the expanded repertoire of GOLPH3 genes in birds could be part of a set of genomic traits that account for their lower susceptibility to cancer than mammals.

### Molecular divergence between duplicated copies of GOLPH3 genes

According to our analyses, the divergence values between duplicated GOLPH3 copies are low. In Galliformes, the divergence values ranged from 1.79% to 4.76%. However, by checking the amino acid alignment, we realized that most of the observed differences are in the first ~50 amino acids of the N-terminal region of the protein (Fig. 3). By estimating amino acid sequence divergence for the N- and C-terminal regions separately, we observed that the divergence values for the C-terminal region ranged from 1.79% to 2.47%, while for the N-terminal region, which represents only ~1/6 of the amino acid sequence, ranged from 15.69% to 17.65%. In this group of birds we found eleven amino acid positions in the alignment that unequivocally distinguish between both paralogs (Fig. 3). Six of them are in the N-terminal region, while the others are in the C-terminal portion of the molecule (Fig. 3). In Anseriformes (Fig. 4), the divergence values range from 1.69% to 4.49%, similar to those estimated for Galliformes. Also, most of the observed differences are in the N-terminal region of the molecule with divergence values ranging from 5.66% to 9.62%. In the case of the C-terminal region, the values varied from 0.8% to 2.9%. In this group of birds there are three amino acid positions that unequivocally distinguish both paralogs (Fig. 4), one of them is located on the N-terminal region of the molecule, whereas the other two in the C-terminal region (Fig. 4). In Neoaves we found the same evolutionary pattern as described for Anseriformes and Galliformes, i.e. most of the amino acid replacements are found in the N-terminal portion of the protein. The divergence values for the whole protein ranged from 1.36% to 3.72%, for the N-terminal region varied from 3.92% to 15.38%, whereas for the C-terminal region went from 0.41% to 2.06%. Unlike the previous cases we did not identify amino acid sites in the alignment that distinguish both paralogs (Fig. 5).

**Figure 4.**
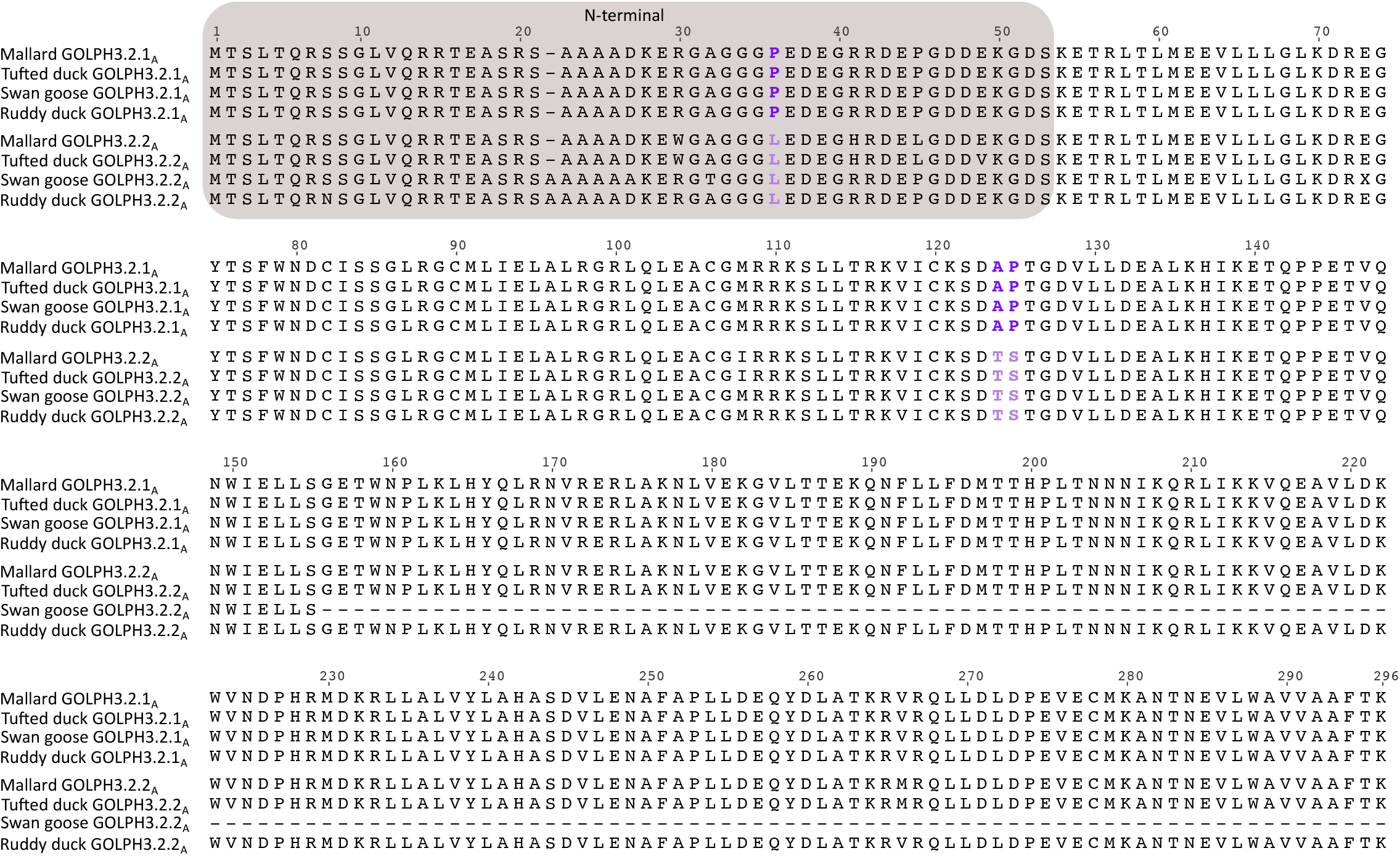
Alignment of Golgi phosphoprotein 3 (GOLPH3) amino acid sequences from mallard (*Anas platyrhynchos*), Tufted duck (*Aythya fuligula*), swan goose (*Anser cygnoides*) and ruddy duck (*Oxyura jamaicensis*). Amino acids positions that differentiate between paralogs are in purple and light purple. The N-terminal region of the protein is shaded.

**Figure 5.**
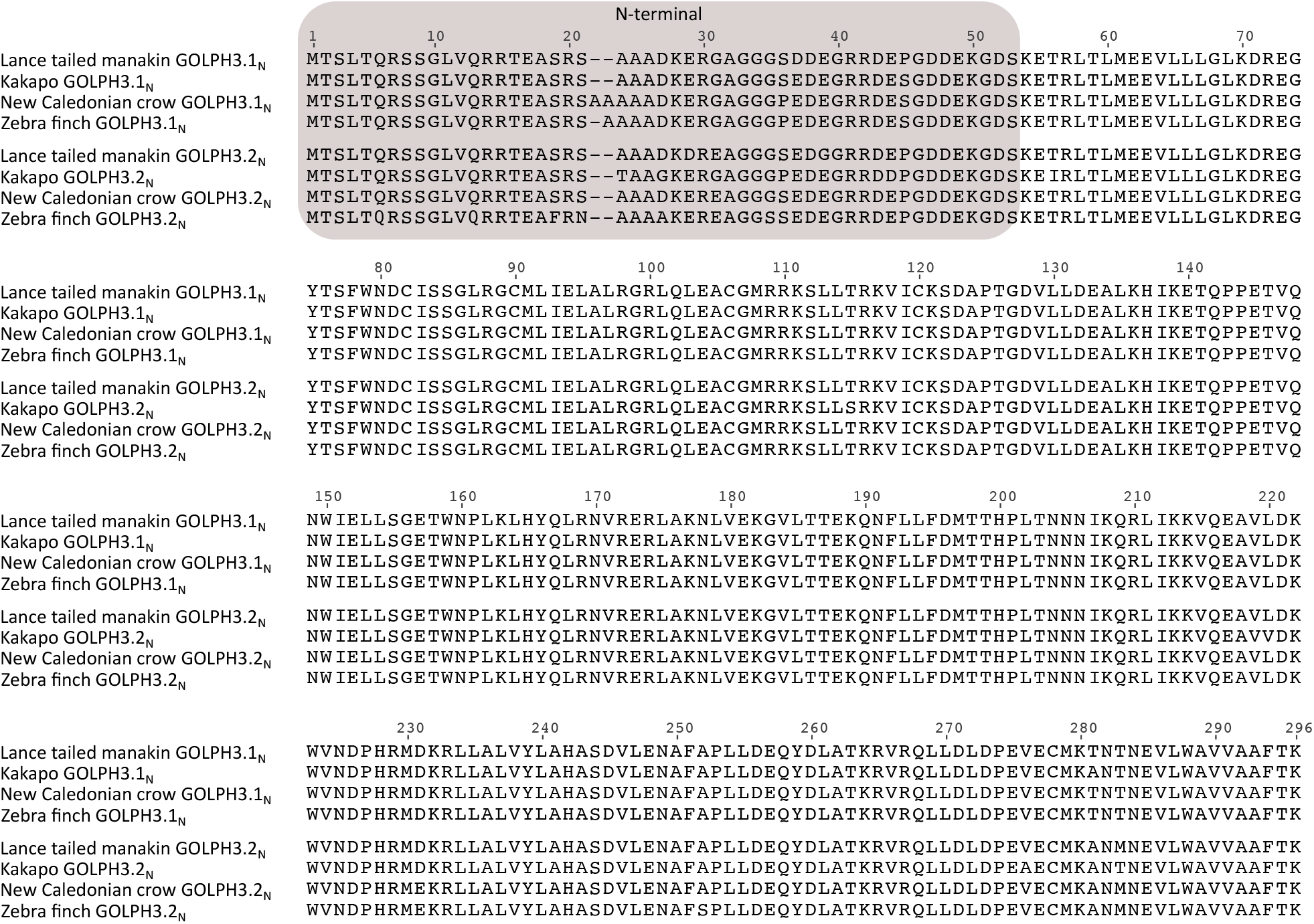
Alignment of Golgi phosphoprotein 3 (GOLPH3) amino acid sequences from lance tailed manakin (*Chiroxiphia lanceolata*), kakapo (*Strigops habroptilus*), New caledonian crow (*Corvus moneduloides*) and zebra finch (*Taeniopygia guttata*). The N-terminal region of the protein is shaded.

These differences in amino acid sequence divergence for the N- and C-terminal regions of the GOLPH3 paralogs could have arisen as a consequence of different structural and functional constraints during its evolution. Secondary structure prediction indicates that the region comprising the first ~40-60 amino acids of GOLPH3 is disordered in a variety of organisms (e.g., yeast, fruit fly, spotted gar, human) that share a common ancestor more than a billion of years ago (Fig. 6A and Supplementary Figure 1). Accordingly, only the crystal structures of N-terminal truncation variants of GOLPH3 and Vps74 (GOLPH3 in yeasts) have been solved ^44,45^. Both structures are remarkably similar (backbone atom root mean square deviation of ~1.0 Å), consisting of a single globular domain that is predominantly α-helical, with a central four-helix bundle surrounded by solvent-exposed loops, and eight amphipathic helices ^44,45^. The overall structure of the N-terminal truncated GOLPH3 protein is unique, with no strong structural homology to known protein folds, resulting so far challenging to predict its function based on its structure. Protein structure homology modeling of GOLPH3.1_GA_ and GOLPH3.2_GA_ of chicken (Supplementary Figure 2) showed that of the divergent amino acids in the C-terminal region only L255 in GOLPH3.1_GA_ and Q256 in GOLPH3.2_GA_ are non-conservative (Fig. 3) ^46^. The position of Q256 is predicted to be exposed at the surface of GOLPH3.2_GA_, like it is for Q260 in human GOLPH3. However, the variant L255 in GOLPH3.1_GA_ is intriguing because the preferred position of leucine residues is buried in regions of proteins facing hydrophobic cores and not exposed on protein surfaces/boundaries such as in this case. None of the divergent amino acids of the C-terminal region in Anseriformes and Neoaves are structurally disfavored. The C-terminal region of GOLPH3 is sufficient for GOLPH3 physical interaction with the membrane of the Golgi apparatus ^1^. This interaction is mediated by a series of highly conserved residues that are postulated to interact with phosphate groups and the inositol ring of phosphatidylinositol 4-phosphate located in the cytosolic leaflet of the Golgi membrane ^44^, set of residues that are also conserved in both copies of GOLPH3 in birds (Figs. 3–5 and Supplementary Figures 1 and 3). In contrast, the N-terminal disordered region has no known function. Intriguingly, some proteins containing disordered regions have the capacity to undergo liquid-liquid phase separation that could result in their partitioning in functional biomolecular condensates also known as membrane-less compartments ^1,47^. However, it is unknown whether GOLPH3 has this capacity. In any case, the distinct amino acid sequence divergence values for the N-terminal disordered region of GOLPH3 suggest a more flexible functional role. Thus, it will be important to determine whether this domain contributes to the functions of GOLPH3 as oncoprotein.

**Figure 6.**
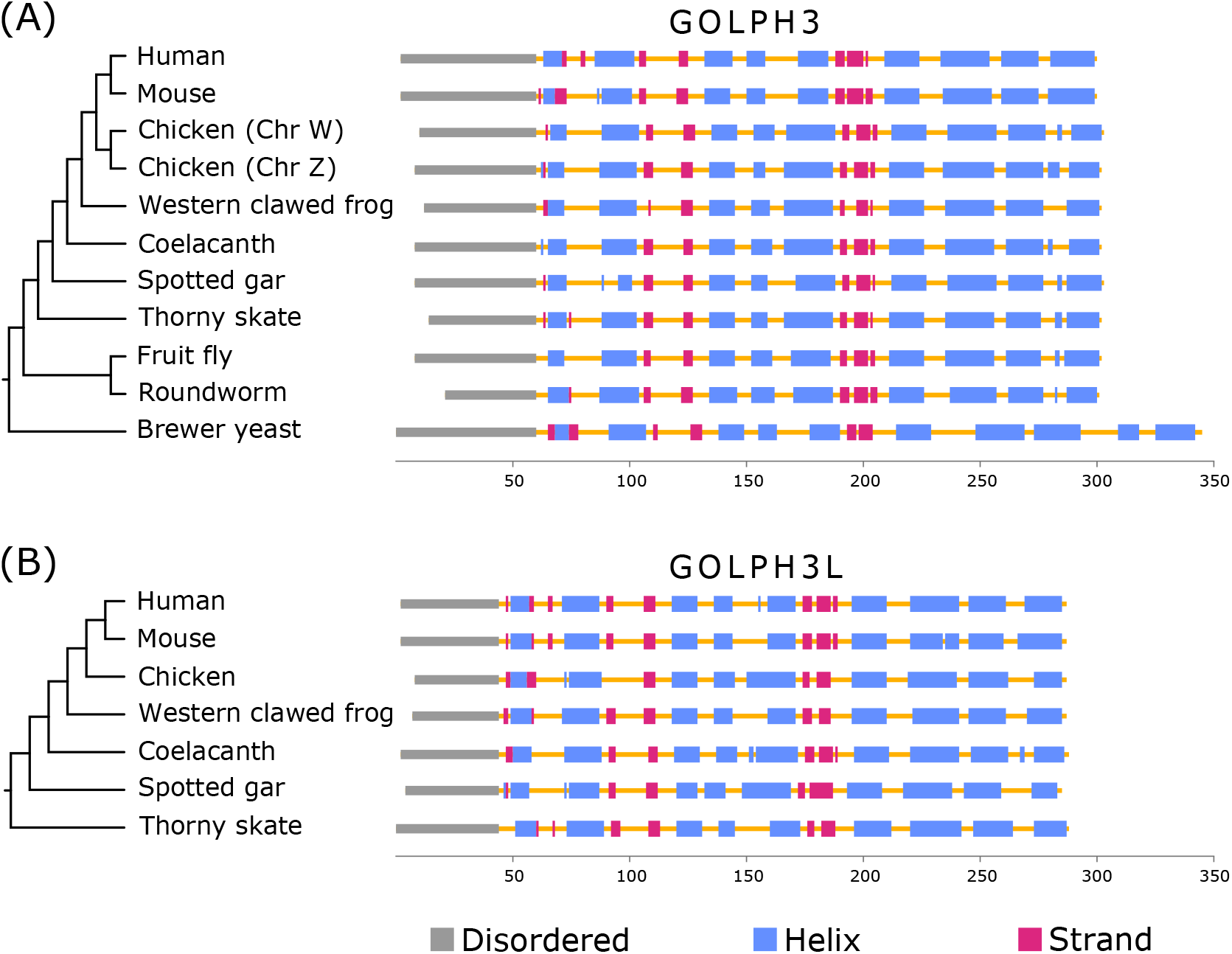
Secondary structure and disordered region predictions of GOLPH3 (A) and GOLPH3L (B) in human (*Homo sapiens*), mouse (*Mus musculus*), chicken (*Gallus gallus*), Western clawed frog (*Xenopus tropicalis*), coelacanth (*Latimeria chalumnae*), spotted gar (*Lepisosteus oculatus*), thorny skate (*Amblyraja radiata*), fruit fly (*Drosophila melanogaster*), roundworm (*Caenorhabditis elegans*) and brewer yeast (*Saccharomyces cerevisiae*).

### Evolution of GOLPH3L paralog

In contrast to GOLPH3, GOLPH3L is largely uncharacterized. Although the amino acid sequences of human GOLPH3 and GOLPH3L are 78% similar (65% identical), it has been suggested that GOLPH3L antagonizes the functions of GOLPH3 ^48^. Despite this, other reports suggest a similar function to GOLPH3 for GOLPH3L in some types of cancer ^49–53^. The evolutionary history of GOLPH3L followed a different trajectory in comparison to the GOLPH3 gene (Fig. 7). In this case, our gene tree recovered the main groups of birds according to the most updated organismal phylogenies ^22–25^; nevertheless, it was not possible to define the relationships among them (Fig. 7). We will assume that the lack of resolution is mainly caused by the limited amount of phylogenetic information contained in a single gene, instead of more complex evolutionary scenarios invoking gene duplications and reciprocal loss in the ancestor of the main groups of birds. Thus, according to our results the GOLPH3L gene was present in the ancestor of birds as a single copy gene (Fig. 2), and this gene was inherited by all descendant lineages (Fig. 2). Thus, GOLPH3L genes in different bird species are 1:1 orthologs. Amino acid divergence values show a similar trend as we described for GOLPH3, i.e., the N-terminal portion of the protein is more divergent than the C-terminal region (Fig. 8). In the case of Galliformes, the divergence values for the N-terminal part of the molecule ranged from 5.71% to 17.14%, while for the C-terminal region it varied from 3.28% to 6.97%. In Anseriformes, the values for the N-terminal region ranged from 7.89% to 26.32%, whereas for the C-terminal portion varied from 2.05% to 4.92%. In the case of Neoves, the evolutionary trend is the same, although the values for the N-terminal region are higher. Thus, the values for the N-terminal part of the molecule ranged from 20% to 43.34%, while for the C-terminal region varied from 4.10% to 17.55%. The secondary structure prediction indicates that this region in GOLPH3L, although shorter than in GOLPH3, is also disordered (Fig. 6B).

**Figure 7.**
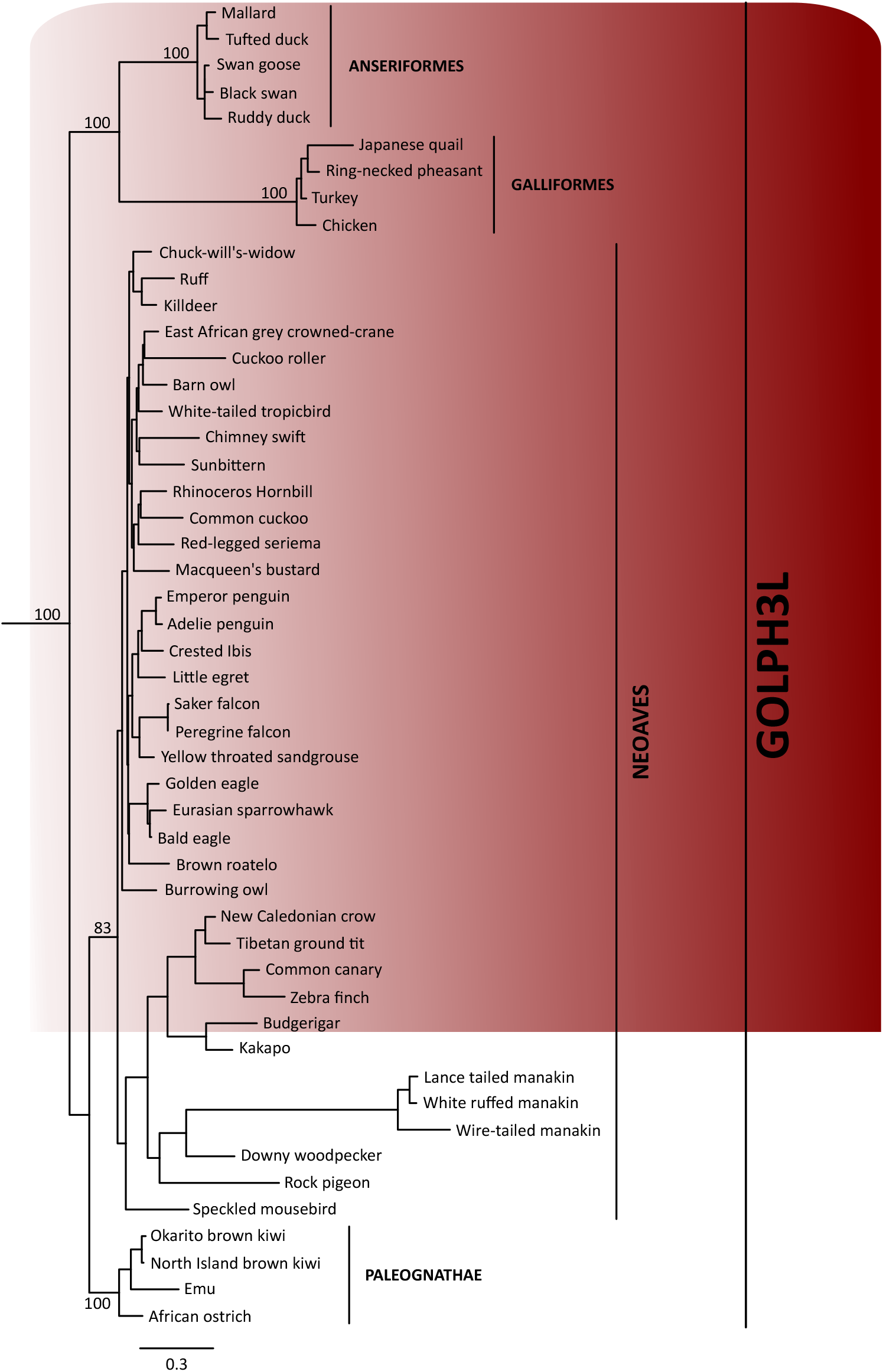
Maximum likelihood tree showing sister group relationships among GOLPH3L genes of birds. Numbers above the nodes correspond to support values from the ultrafast bootstrap routine. GOLPH3L sequences from crocodiles and turtles were used as outgroups (not shown). The scale denotes substitutions per site.

**Figure 8.**
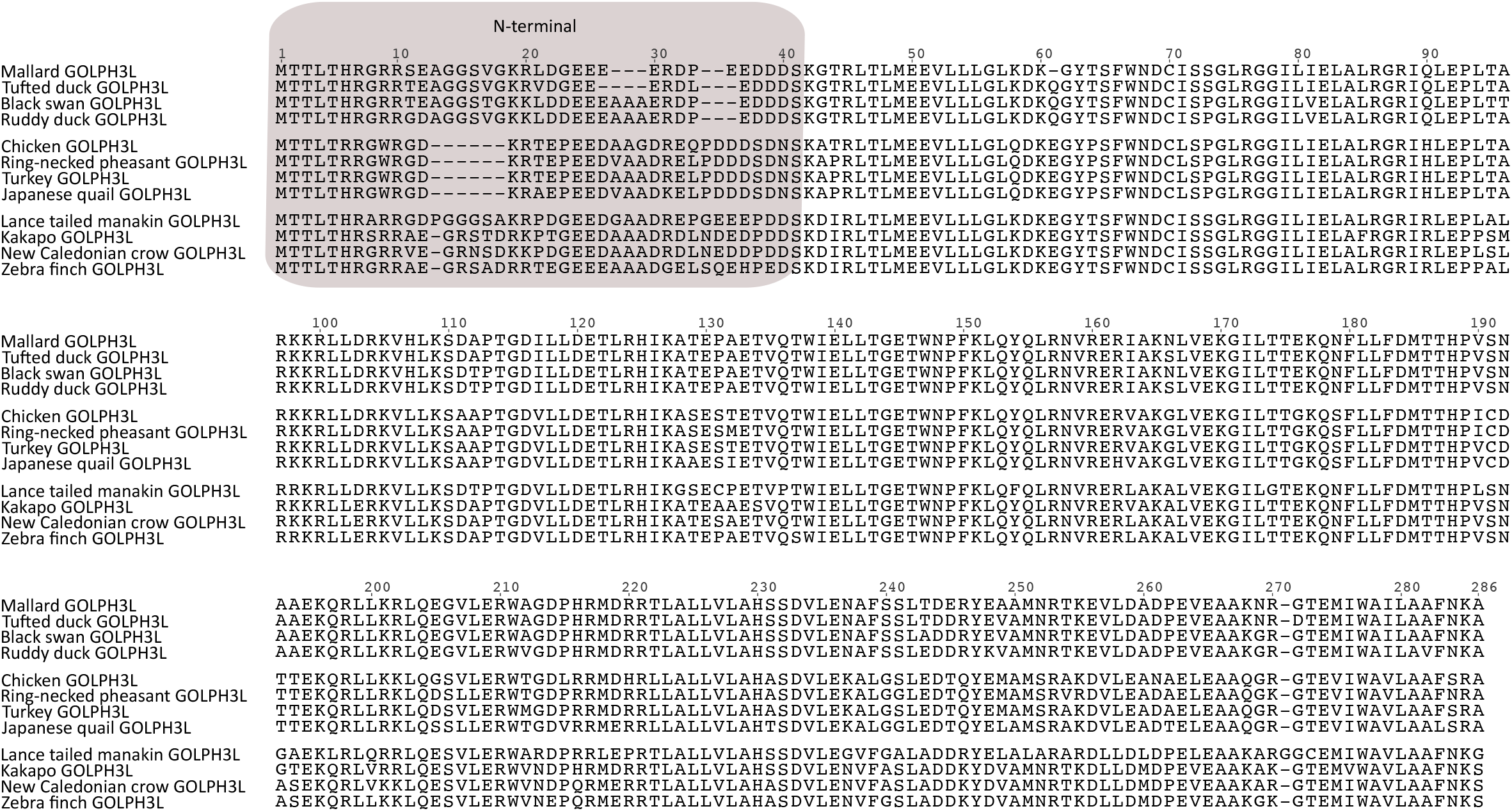
Alignment of Golgi phosphoprotein 3L (GOLPH3L) amino acid sequences from mallard (*Anas platyrhynchos*), Tufted duck (*Aythya fuligula*), black swan (*Cygnus atratus*), ruddy duck (*Oxyura jamaicensis*), chicken (*Gallus gallus*), ring-necked pheasant (*Phasianus colchicus*), turkey (*Meleagris gallopavo*), japanese quail (*Coturnix japonica*), lance tailed manakin (*Chiroxiphia lanceolata*), kakapo (*Strigops habroptilus*), New caledonian crow (*Corvus moneduloides*) and zebra finch (*Taeniopygia guttata*). The N-terminal region of the protein is shaded.

One thing that seems interesting is the number of changes accumulated in the branch leading to Galliformes and to manakins (Fig. 7). This phenomenon could be indicative of an acceleration of the rate of fixation of amino acid changing mutations in the ancestors of both groups. To test this hypothesis, we estimated the omega value (d_N_/d_S_), i.e., the ratio of non-synonymous (d_N_) to synonymous substitutions (d_S_), in the branches leading to both groups. In brief, if non-synonymous substitutions are neutral, then the rate of fixation of d_N_ and d_S_ will be very similar, and d_N_/d_S_ ≈ `l. Under negative selection, most non-synonymous substitutions are deleterious, and d`_N_/d_S_ < 1. Finally, under positive selection non-synonymous (d_N_) replacements are advantageous and will be fixed at a greater rate than synonymous substitutions (d_S_) and in consequence d_N_/d_S_ > 1 ^54^. According to our analyses, in the ancestor of Galliformes the model in which the omega value was estimated from the data was not significantly different from the model in which the omega value was fixed to 1 (neutral evolution)(LRT=0.142, P > 0.05). On the other hand, in the case of manakins the model in which the omega value was estimated from the data (d_N_/d_S_=6.2) was significantly different from the null hypothesis of neutral evolution (LRT=7.19, P < 0.01), indicating that the rate of fixation of non-synonymous substitutions (d_N_) is higher in comparison to the neutral expectation (d_S_) and suggesting an event of positive selection in the ancestor of manakins. According to the Bayes Empirical Bayes (BEB) approach five sites (152C/G, 191R, 230A, 233R and 263G) were inferred under positive selection with a posterior probability higher than 0.95. All of them are located in the C-terminal region of the protein. Given the limited understanding of the biological functions associated with the GOLPH3 gene family, in particular of the GOLPH3L gene ^3^, it is challenging to explain the consequences of positive selection in GOLPH3L in this group of birds. However, it could be interesting to carry out functional assays in which the performance of manakins GOLPH3L protein is compared to the one in other birds.

### Expression pattern of GOLPH3 gene family members

Our next step was to investigate the expression pattern of the GOLPH3 gene family members, especially for the duplicated copies derived independently in different groups of birds. To do this, we mapped RNASeq reads to reference gene sequences in the chicken (*Gallus gallus*) and mallard (*Anas platyrhynchos*) and examined transcript abundance in a panel of nine tissues (Fig. 9). It is important to say that in chicken, the duplicated GOLPH3 copies are located on chromosome Z (GOLPH3.2_GA_) and W (GOLPH3.1_GA_), while GOLPH3L is on chromosome 25. Therefore, females (ZW) can express all gene family members, whereas males (ZZ) can only express GOLPH3.2_GA_ and GOLPH3L. This situation is somewhat similar to the allelic trichromacy observed in New World monkeys, where some females (XX) possess trichromatic color vision due to a polymorphism of an opsin gene located on chromosome X, while males (XY) are all dichromatic ^55^. The case is different in mallard, as one copy is located on chromosome Z (GOLPH3.2.1_A_) but GOLPH3.2.2_A_ and GOLPH3L are autosomal genes, so both sexes can potentially express all paralogs.

**Figure 9.**
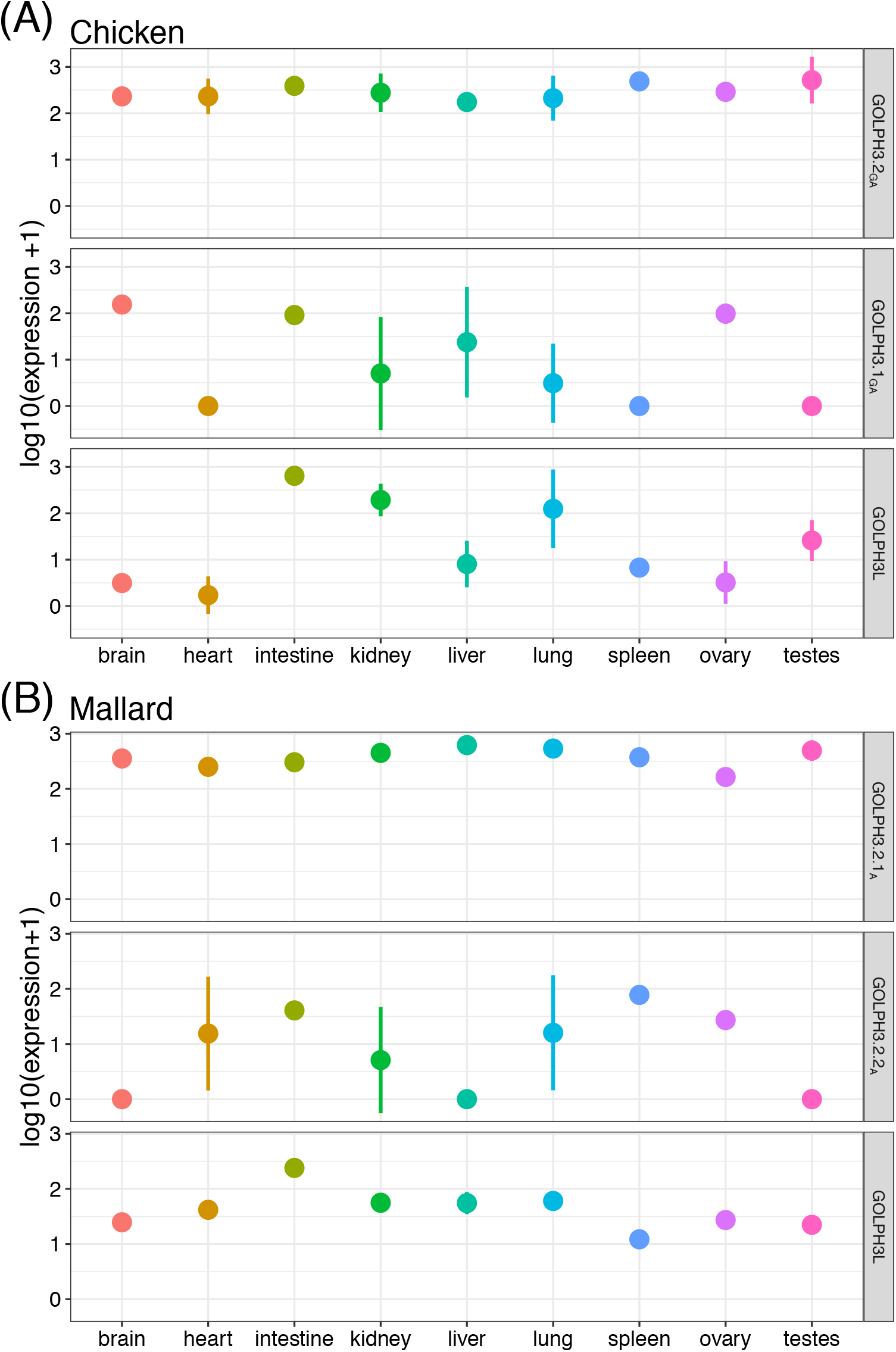
Transcript abundance measurements of GOLPH3 paralogs across a panel of nine tissues in A) chicken (*Gallus gallus*) and B) mallard (*Anas platyrhynchos*). The mean and standard deviation are plotted for three replicates per tissue. In many cases the standard deviation was smaller than the size of the circle representing the mean and appears hidden.

In both species, the paralog located on the Z chromosome (GOLPH3.2_GA_ in chicken and GOLPH3.2.1_A_ in mallard) was highly and ubiquitously expressed across all tissues (Fig. 9). We collected chicken libraries from both male and female tissues, and unfortunately the sex of the individuals for some tissues was not declared (Supplemental Table S2). As such GOLPH3.1_GA_ exhibited variable expression from mixed sex sampling (Fig 9A). Thus, GOLPH3.1_GA_ was highly expressed in the brain and ovary where all libraries were constructed from female individuals (Fig 9A). Although we do not know the sex of the individuals for the intestine libraries of the chicken, based on the consistent expression of the gene located on chromosome W, we can presume that they were all from female individuals (Fig. 9A). As a validation of what we mentioned above, GOLPH3.1_GA_ was not expressed in all known male tissue libraries, which was most noticeable in the male specific testes (Fig. 9A). All autosomal paralogs in both species were variably expressed among and within tissues (Fig 9). In the case of the chicken GOLPH3L, we recovered expression in all tissues but at variable levels, from low values in the brain and heart to higher values in the intestine and kidney (Fig. 9A). By contrast GOLPH3L was universally expressed in all mallard tissues (Fig. 9B). GOLPH3.2.2_A_ in mallard was not expressed in the brain, liver and testes, but highly expressed in the ovary, spleen and intestine. In both species, all GOLPH3 paralogs were consistently expressed in the intestine (Fig. 9).

In humans, both paralogs are ubiquitously expressed across all tissues ^56^, suggesting that they are required for the maintenance of basic cellular functions ^57^; however, GOLPH3L seems to be expressed at lower levels. Similarly, the relative expression levels of GOLPH3 and GOLPH3L in several mammalian cell lines with epithelial, fibroblast, myeloid and neuronal characteristics, and in a variety of tissues from mice also indicates that GOLPH3 is also ubiquitously expressed at higher levels than GOLPH3L ^48^. Further, GOLPH3L is expressed more in cells with secretory epithelial characteristics ^48^, suggesting a distinct function for this gene family member. The expression pattern observed in the mallard is similar to what is observed in model species (Fig. 9B). One of the GOLPH3 duplicates (GOLPH3.2.1_A_) is expressed in all examined tissues at high levels, while GOLPH3L is also expressed in all tissues, but at lower levels. The expression of the other duplicate (GOLPH3.2.2_A_) is variable, including tissues in which it is not detected (Fig. 9). The case of the chicken seems to be more dissimilar. In this species one of the GOLPH3 duplicates (GOLPH3.2_GA_) possesses an expression pattern similar to the human GOLPH3 (Fig. 9A), however the other two copies seem to follow a specific expression pattern (Fig. 9A).

### Conclusions

Our study shows that the evolution of the GOLPH3 gene family followed a more complicated evolutionary pathway than previously thought. Although the history of the GOLPH3L paralog is according to the current knowledge, the one of GOLPH3 is not. The most exciting thing about the evolution of GOLPH3 in birds is that they possess extra GOLPH3 gene copies never described before, and that all main groups independently originated their repertoire. In other words, they do not have the same evolutionary origin, and in consequence, they are not 1:1 orthologs and are not directly comparable. Most paleognaths retained the ancestral condition of a single gene copy, whereas Galliformes, Anseriformes, and Neoaves possess duplicated copies that were originated independently. Thus, birds represent a natural experiment of gene copy number variation ^58^, that in addition to the differences in expression of individuals of different sex, could help us improve our understanding of the biological functions associated with the GOLPH3 gene family. Our results also highlight the power of manually curating genetic data to define gene repertoires, and the reconciliation of gene trees with species trees ^59^ to understand the duplicative history of gene families to perform biologically meaningful comparisons ^21,60^. Finally, the conservation of the N-terminal portion of GOLPH3 paralogs as a disordered region for more than a billion of years of evolution and the fact that it displays a higher degree of divergence among species, compared to the C-terminal portion, strongly suggests that it performs an essential, specialized and adapted cellular function conserved in distantly related species like yeasts and humans that remains to be elucidated.

## Material and Methods

### DNA sequences and phylogenetic analyses

We performed searches for GOLPH3 sequences in avian genomes in the National Center for Biotechnology Information (NCBI) ^61^ and the Ensembl v.102 databases ^62^. We retrieved orthologs and paralogs from the NCBI ^61^ using the chicken (*Gallus gallus*), zebra finch (*Taeniopygia guttata*), and mallard (*Anas platyrhynchos*) sequences using the program blast (blastn) ^63^ against the non-redundant database (nr) with default parameters. Additionally, we also retrieved sequences from the Ensembl v.102 database ^62^. In cases where sequences are not complete we manually annotated them. To do so, we first identified the genomic fragment containing the GOLPH3 gene in Ensembl v.102 ^62^ or NCBI databases ^64^. Once identified, genomic fragments were extracted, including flanking genes. After extraction, we manually annotated GOLPH3 genes by comparing known exon sequences from a species that share a common ancestor most recently in time to the species of which the genomic piece is being annotated using the program Blast2seq v2.5 ^65^ with default parameters. Accession numbers and details about the taxonomic sampling are available in Supplementary Table S1.

We performed separate phylogenetic analyses for GOLPH3 and GOLPH3L paralogs. Amino acid sequences were aligned using MAFFT v.7 ^66^, allowing the program to choose the alignment strategy (L-INS-i in both cases). Nucleotide alignments were generated using the amino acid alignments as templates using the software PAL2NAL ^67^. We used the proposed model tool of IQ-Tree v.1.6.12 ^68^ to select the best-fitting model of codon substitution, which selected MGK+F1X4+R3 for GOLPH3 and MGK+F3X4+G4 for GOLPH3L. This approach uses a more realistic description of the evolutionary process at the protein-coding sequence level by incorporating the genetic code structure in the model. We used the maximum likelihood method to obtain the best trees using the program IQ-Tree v1.6.12 ^69^. We assessed support for the nodes using three strategies: a Bayesian-like transformation of aLRT (aBayes test) ^70^, SH-like approximate likelihood ratio test (SH-aLRT) ^71^ and the ultrafast bootstrap approximation ^72^. In each case (GOLPH3 and GOLPH3L), we carried out 25 independent runs to explore the tree space, and the tree with the highest likelihood score was chosen. In both cases, GOLPH3 and GOLPH3L sequences from crocodiles and turtles were used as outgroups (Supplementary Table S1).

### Molecular Evolution analysis

To measure variation in functional constraint among the GOLPH3L genes and to test for evidence of positive selection, we estimated the omega parameter (d_N_/d_S_), using a maximum-likelihood approach ^73^ implemented in the CODEML module of the program PAML v.4.8a ^74^. We implemented branch-site models, which explore changes in the omega parameter for a set of sites in a specific branch of the tree to assess changes in their selective regime ^75^. In this case, we conducted two separate analyses. In the first, the ancestral branch of Galliformes was labeled as the foreground branch, while in the second, the branch leading to manakins was labeled as a foreground branch. We compared the modified model A ^75–77^, in which some sites are allowed to change to an omega value > 1 in the foreground branch, with the corresponding null hypothesis of neutral evolution using a Likelihood Ratio Test (LRT). Three starting omega values (0.5, 1, and 2) were used to check the existence of multiple local optima. The Bayes Empirical Bayes (BEB) method was used to identify sites under positive selection ^78,79^.

### Secondary structure and disordered region prediction and protein structure homology modeling

Secondary structure and disordered region prediction was performed using the PredictProtein server (https://predictprotein.org/)^80^. Multiple sequence alignment was performed using MAFFT v.7 server (https://mafft.cbrc.jp/alignment/server/) with default parameters ^81^. Multiple sequence alignment editing was performed using Jalview software v.2.11.1.3 ^82^. Protein structure homology modeling was performed using the SWISS-MODEL server (https://swissmodel.expasy.org/)^83^. Structural figures were prepared with PyMOL Molecular Graphics System, Version 2.0.6 Schrödinger, LLC.

### Transcript abundance analyses

GOLPH3 transcript abundance was measured in chicken (*Gallus gallus*) and mallard (*Anas platyrhynchos*). We collected three RNASeq libraries from brain, heart, intestine, kidney, liver, lung, spleen, ovary, and testis from each species gathered from the NCBI Short Read Archive (SRA) ^84^. Accession numbers can be found in Supplemental Table S2. Reference transcript sequences were collected from Ensembl v.102 ^62^, and we only included the longest transcript for each gene. For each library, adapters were removed using Trimmomatic 0.38 ^85^, and reads were filtered for quality using the parameters HEADCROP:5, SLIDINGWINDOW:5:30, and MINLEN:50. We mapped quality filtered paired-end RNAseq reads back to reference sequences using Bowtie 1.2.2 ^86^ and default parameters of RSEM ^87^. Normalization of raw read counts for each species was performed using the estimateSizeFactors and estimateDispersions functions in DESeq2 v1.26 ^88^.

## Acknowledgements

This work was supported by Fondo Nacional de Desarrollo Científico y Tecnológico from Chile (FONDECYT 1210471) and Millennium Nucleus of Ion Channels Associated Diseases (MiNICAD), Iniciativa Científica Milenio, Ministry of Economy, Development and Tourism from Chile to JCO, the US Dept. of Education HSI-STEM grant P031C110114-15 to MWV, Fondo Nacional de Desarrollo Científico y Tecnológico from Chile (FONDECYT 1180957) to FJM and LVC, Fondo Nacional de Desarrollo Científico y Tecnológico from Chile (FONDECYT 1211481) to GAM and Vicerrectoría de Investigación, Desarrollo y Creación Artística of Universidad Austral de Chile (JCO, FJM, LVC, GAM).

## Author contributions

JCO and GAM designed the study. MWV, JG, KZ, JCO, GAM collected and/or analyzed data. JCO and GAM wrote the manuscript. MWV, LV-C, FJM reviewed and edited the manuscript. All authors contributed to the article and approved the submitted version.

